# Identifying EEG Biomarkers of Depression with Novel Explainable Deep Learning Architectures

**DOI:** 10.1101/2024.03.19.585728

**Authors:** Charles A. Ellis, Martina Lapera Sancho, Robyn L. Miller, Vince D. Calhoun

## Abstract

Deep learning methods are increasingly being applied to raw electro-encephalogram (EEG) data. However, if these models are to be used in clinical or research contexts, methods to explain them must be developed, and if these models are to be used in research contexts, methods for combining explanations across large numbers of models must be developed to counteract the inherent randomness of existing training approaches. Model visualization-based explainability methods for EEG involve structuring a model architecture such that its extracted features can be characterized and have the potential to offer highly useful insights into the patterns that they uncover. Nevertheless, model visualization-based explainability methods have been underexplored within the context of multichannel EEG, and methods to combine their explanations across folds have not yet been developed. In this study, we present two novel convolutional neural network-based architectures and apply them for automated major depressive disorder diagnosis. Our models obtain slightly lower classification performance than a baseline architecture. However, across 50 training folds, they find that individuals with MDD exhibit higher β power, potentially higher δ power, and higher brain-wide correlation that is most strongly represented within the right hemisphere. This study provides multiple key insights into MDD and represents a significant step forward for the domain of explainable deep learning applied to raw EEG. We hope that it will inspire future efforts that will eventually enable the development of explainable EEG deep learning models that can contribute both to clinical care and novel medical research discoveries.

## 1 Introduction

While deep learning methods are often applied to both raw resting-state electroencephalography (EEG) data [1–17] and extracted EEG features [18–33] in research contexts for neuropsychiatric disorder diagnosis, they are increasingly being applied to raw data with the goal of developing more flexible models that can use all of the information in available data, rather than just a subset of that information [1–17]. Nevertheless, compared to traditional machine learning methods or deep learning methods applied to extracted features [18–33], the training of deep learning models on raw resting-state EEG presents greater challenges related to explainability. As such, in recent years, a growing number of studies have begun to present explainability methods for raw EEG. Some of those methods have focused on post-hoc analyses [1, 2, 10–17], and some have focused on the development of novel architectures that offer enhanced interpretability and enable the development of more reliable post-hoc analyses [3–10]. In this study, we present two new convolutional neural network (CNN) architectures that afford greater interpretability, comparing them to a baseline architecture in terms of model performance and traditional post hoc spatial and spectral explainability, and further present two new post hoc analyses uniquely enabled by our architectures that provide disorder-related insights into the features extracted by our models. We apply the architectures for multichannel EEG-based diagnosis of major depressive disorder (MDD), and analyze them across a large number of folds to identify generalizable EEG biomarkers of MDD.

While machine learning methods have been applied to resting-state EEG data for at least the past two decades [27, 28], the rise of deep learning in the past decade has resulted in increased application of machine learning and deep learning methods to EEG. Many of these studies have used manually engineered, extracted EEG features (e.g., spectral power [26–33] or connectivity [20–25]). Nevertheless, the use of manually engineered EEG features inherently limits the amount of information available for use by machine learning models. Moreover, deep learning architectures perform automated feature extraction that has the potential to uncover the most salient information in raw data [34]. The combination of the potentially richer feature space in raw EEG data and the potential for deep learning methods to uncover the most salient features has occasioned the application of deep learning models to raw EEG data with the goal of maximizing the utility of learned features [34]. Nevertheless, deep learning methods applied to raw EEG data tend to be less explainable than methods applied to extracted features, which is highly problematic both for extracting scientific insights (i.e., biomarkers) from deep learning models [35] and for healthcare applications [36].

Traditional machine learning approaches and deep learning approaches applied to manually engineered features tend to be more explainable because of the relatively large number of methods developed in other applications. For example, methods like elastic net [18] are inherently interpretable, and methods like permutation importance [19] can easily be applied to traditional machine learning models. Dozens of other approaches have been developed for tabular data in traditional machine learning models [37–39]. Furthermore, deep learning models trained on manually engineered features can be combined with any of a variety of explainability methods [40–42]. In contrast, the explanation of deep learning models trained on non-stationary raw resting-state EEG data is a non-trivial matter. While identifying key electrodes only requires a relatively straightforward adaptation of existing ablation or gradient-based explainability approaches [43], identifying frequency bands, waveforms, or spatial interactions of global importance requires more specialized, domain-specific approaches.

As such, a growing number of methods have been developed to explain deep learning models trained on raw, resting-state EEG data. These methods can be categorized based on the type of EEG features into which the provide insight. For example, they can provide insight into key frequency bands (spectral) [11–14], waveforms (temporal) [3–5], channels (spatial) [2, 12, 43–46], or spatially correlated activity (interaction) [17, 47]. They can also be categorized based upon the broader type of explainability method from which they are adapted (e.g., activation maximization [1, 2, 10], data perturbation [11–14], gradient-based feature attribution [15–17], model visualization [3–5, 10], activation analysis [6, 7, 10], or interpretable models [8, 9]). Nevertheless, while existing methods provide helpful insights, they have some noteworthy limitations.

For example, activation maximization methods can be computationally intensive [1], generally operate best with samples of very short lengths [48, 49], and have not yet been extended from single-channel to multichannel EEG data. Data perturbation methods have the potential to create out-of-distribution samples that can reduce the faithfulness of explanations [39]. Gradient-based feature attribution methods [50] have been shown to sometimes yield questionable explanations in other domains [51], and intrinsically interpretable models (i.e., having specialized first layer filters that only extract a specific frequency band) limit the available feature space, which limits the utility of training with raw EEG data [8, 9]. Among all these approaches, model visualization-based approaches offer the potential for highest quality explanations without limiting the available feature space. Model visualization-based approaches generally involve CNNs with long first-layer filters that can be visualized and characterized [3–5, 10]. However, they do have some limitations. Namely, they are often shallower networks with slightly lower levels of performance [4, 5]. Moreover, most model visualization studies do not involve multi-channel EEG, and to date, have not been adapted for insight into spatial interactions. Additionally, most explainable deep learning studies for raw EEG have another key drawback.

Most studies make methodological contributions [1, 4–6, 11, 13, 45, 52] or simply explain a new architecture [2, 3, 7–10]. Their goal is not necessarily to identify reliable EEG biomarkers. This is problematic if explainable deep learning approaches are ever to be used for identifying biomarkers of neuropsychiatric disorders, because deep learning models can learn a near-infinite distribution of features and decision boundaries [53], many of which will have comparable performance on a given task. The features and decision boundaries learned by a model are dependent upon its initialization and architectural details and the random distribution of samples across the training, validation, and test sets. Thus, when many analyses only consider a single model fold [1–9, 14] or a small number of folds [10–13, 45, 54], the overall reliability or generalizability of their findings from a scientific perspective is likely limited. As such, if deep learning models are ever to be leveraged for novel insights into EEG, the field must develop explainability approaches that can be integrated across large numbers of models [55].

In this study, we present two novel CNN architectures for multi-channel EEG-based MDD diagnosis. The two architectures are among the first to enable multi-channel model visualization-based explainability analyses and are uniquely structured to provide novel insights into the interaction of features learned by models across channels. We compare the two architectures to a baseline CNN architecture that has been used in previous MDD classification studies in terms of model performance and spatial and spectral explainability with two pre-existing post hoc explainability analyses. We then perform a series of novel model visualization and activation-based explainability analyses that are uniquely enabled by our architectures. We perform all explainability analyses across 50 folds and demonstrate how explanations can be combined across all folds to obtain insights into EEG biomarkers of MDD that are corroborated by existing research. We hope that our novel architectures and subsequent explainability analyses will find use in future studies and that our study will provide useful guidance for future studies seeking to gain novel insights through the application of explainable deep learning methods to raw resting-state EEG.

## 2 Methods

In this section, we describe and discuss our methodology. (1) We use a multichannel resting state EEG dataset with individuals with MDD (MDDs) and healthy controls (HCs). (2) Train a pre-existing baseline architecture (M1) and 2 novel architectures that enable model visualization-based explainability (M2 and M3). (3) We compare the performance and patterns learned (spatial and spectral explainability) by the 3 architectures. (3) We cluster and characterize the first layer filters extracted by M2 and M3 across folds. (4) Lastly, we perform a pair of novel activation analyses that compare how activations for each cluster of filters in each channel and how correlations between activations for each channel differ between MDDs and HCs.

### 2.1 Description of Dataset

We used a resting state EEG dataset composed of 5-10 minute recordings from 30 MDDs and 28 HC [56]. The dataset has been used in multiple deep learning studies [17, 47, 57, 58]. Recordings were performed with eyes closed, at a sampling rate of 256 Hertz (Hz), and in the standard 10-20 format with 64 electrodes. As in previous studies on automated EEG diagnosis of neuropsychiatric disorders [17, 47, 58–60], we used 19 channels: Fp1, Fp2, F7, F3, Fz, F4, F8, T3, C3, Cz, C4, T4, T5, P3, Pz, P4, T6, O1, and O2. Moreover, because previous studies have shown that deep learning models can obtain high levels of performance on our dataset with a lower sampling rate [58], we decimated the data to 100 Hz. Upon downsampling, we separately z-score normalized each channel of each recording to make model training easier. We lastly segmented the data into 30-second epochs with a sliding window approach. We used a 2.5-second step size to effectually augment the number of available samples for training. Our final dataset contained 2,840 HC samples and 2,850 MDD samples.

### 2.2 Description of Model Development

We developed 2 new CNN architectures and compared them to an existing CNN that has been used in previous studies involving neuropsychiatric disorder classification [58]. In this section, we first describe each of the 3 architectures before describing our overall training approach. We used identical training procedures for each architecture. All code is publicly available at: https://github.com/cae67/InterpretableConv.

#### Model 1 (M1) – Preexisting Architecture

We adapted a baseline architecture that has been used in a variety of previous studies involving neuropsychiatric disorder diagnosis. It was first developed for schizophrenia classification [12, 54, 59] but has since been used in for MDD classification [17, 47, 58]. Table 1 shows our model architecture.

**Table 1.**
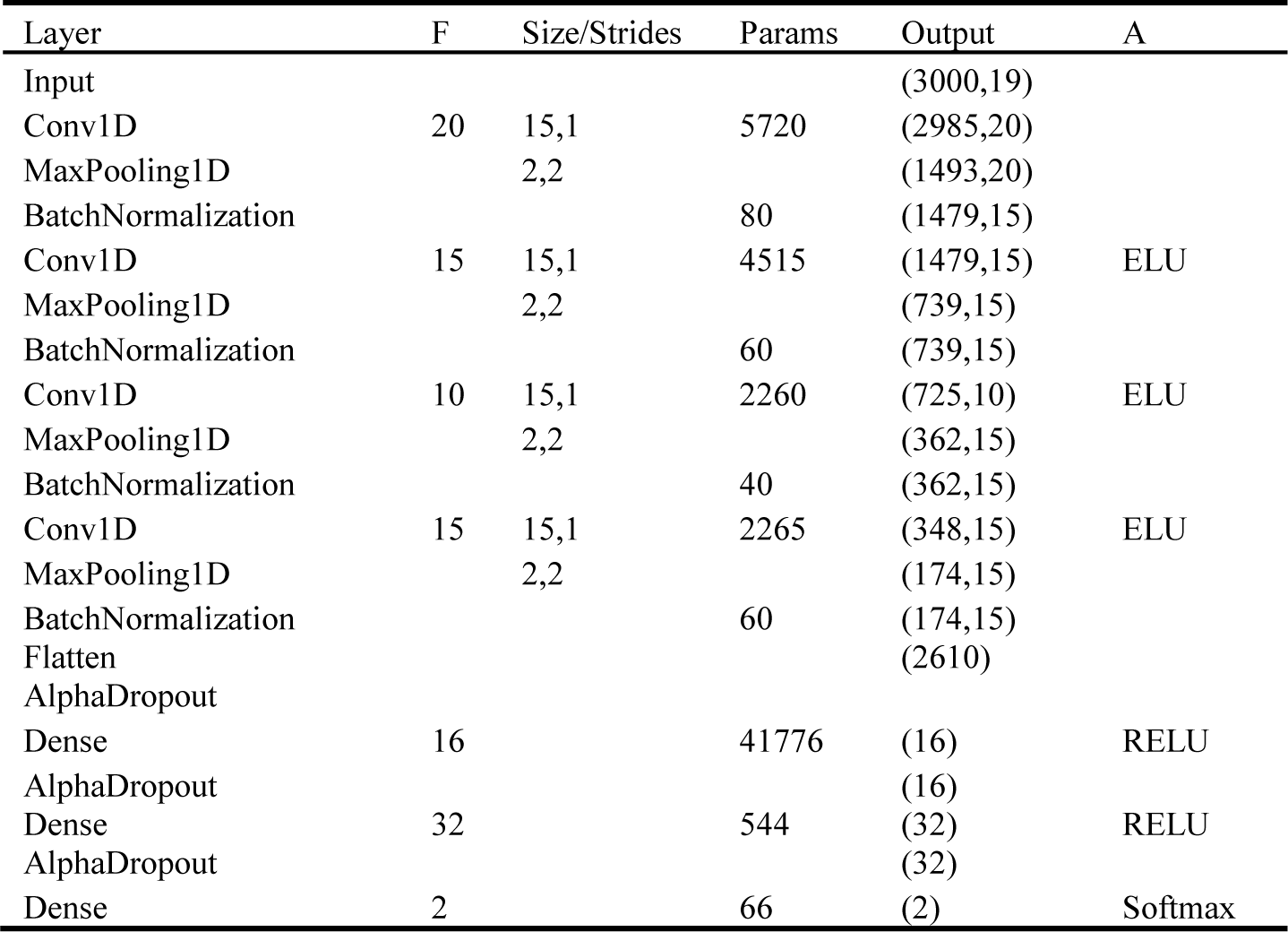
M1 Architecture, where F = number of filters or nodes, A = Activations. All conv layers use padding=’valid’, the he_normal initializer, and a max norm kernel constraint with a maximum value of 1. A learning rate of 5e-5 was used. The first AlphaDropout Layer has a rate of 0.3, and the remaining AlphaDropout Layers have rates of 0.5. The total number of parameters is 57,386, with 57,266 trainable parameters. Layer names are from the Keras package [61].

| Layer | F | Size/Strides | Params | Output | A |
| --- | --- | --- | --- | --- | --- |
| Input |  |  |  | (3000,19) |  |
| Conv1D | 20 | 15,1 | 5720 | (2985,20) |  |
| MaxPooling1D |  | 2,2 |  | (1493,20) |  |
| BatchNormalization |  |  | 80 | (1479,15) |  |
| Conv1D | 15 | 15,1 | 4515 | (1479,15) | ELU |
| MaxPooling1D |  | 2,2 |  | (739,15) |  |
| BatchNormalization |  |  | 60 | (739,15) |  |
| Conv1D | 10 | 15,1 | 2260 | (725,10) | ELU |
| MaxPooling1D |  | 2,2 |  | (362,15) |  |
| BatchNormalization |  |  | 40 | (362,15) |  |
| Conv1D | 15 | 15,1 | 2265 | (348,15) | ELU |
| MaxPooling1D |  | 2,2 |  | (174,15) |  |
| BatchNormalization |  |  | 60 | (174,15) |  |
| Flatten |  |  |  | (2610) |  |
| AlphaDropout |  |  |  |  |  |
| Dense | 16 |  | 41776 | (16) | RELU |
| AlphaDropout |  |  |  | (16) |  |
| Dense | 32 |  | 544 | (32) | RELU |
| AlphaDropout |  |  |  | (32) |  |
| Dense | 2 |  | 66 | (2) | Softmax |

#### Models 2 and 3 (M2 and M3) – Novel Architecture

We develop a pair of novel CNN architectures inspired by an architecture for single channel sleep stage classification that was first presented in [10] and later adapted for model visualization-based explainability [4–6]. As shown in Tables 2 and 3, M2 and M3, respectively, first take a sample as an input, concatenate each channel along the time dimension, and apply a 1-dimensional convolutional layer with a filter length of 50 points (i.e., ½ of the sampling rate or 0.5 seconds). Afterwards, a 2-dimensional convolutional layer is applied to consolidate information in each channel (i.e., consolidating all filter activations in a channel to a single time-series), and global average pooling is applied to obtain 1 feature value per channel. For M2, the features are then fed into a nonlinear classifier (i.e, a dense layer with a ReLU activation function folloed by a dense layer with a softmax activation function), and for M3, the resulting features are fed into linear classifier (i.e., a dense layer with a softmax activation function).

**Table 2.** M2 Architecture, where F = number of filters or nodes, A = Activations. All conv layers use padding=’valid’, the he_normal initializer, and a max norm kernel constraint with a maximum value of 1. A learning rate of 0.001 was used. The total number of parameters is 1,643, with 1,565 trainable parameters. Layer names are from the Keras package [61].

| Layer | F | Size/Strides | Params | Output | A |
| --- | --- | --- | --- | --- | --- |
| Input |  |  |  | (3000,19) |  |
| Transpose |  |  |  | (19,3000) |  |
| Reshape |  |  |  | (57000,1) |  |
| Conv1D | 20 | 50,50 | 1020 | (1140,20) | ELU |
| Expand_dims |  |  |  | (1140,20,1) |  |
| Split |  |  |  | 19 (60,20,1) |  |
| Concatenate |  |  |  | (60,20,19) |  |
| Transpose |  |  |  | (60,19,20) |  |
| BatchNormalization |  |  | 80 | (60,19,20) |  |
| Conv2D | 1 | 10,10 | 201 | (6,19,1) | ELU |
| Squeeze |  |  |  | (6,19) |  |
| GlobalAveragePooling1D |  |  |  | (19) |  |
| BatchNormalization |  |  | 76 | (19) |  |
| Dense | 12 |  | 240 | (12) | RELU |
| Dropout |  |  |  | (12) |  |
| Dense | 2 |  | 26 | (2) | Softmax |

**Table 3.**
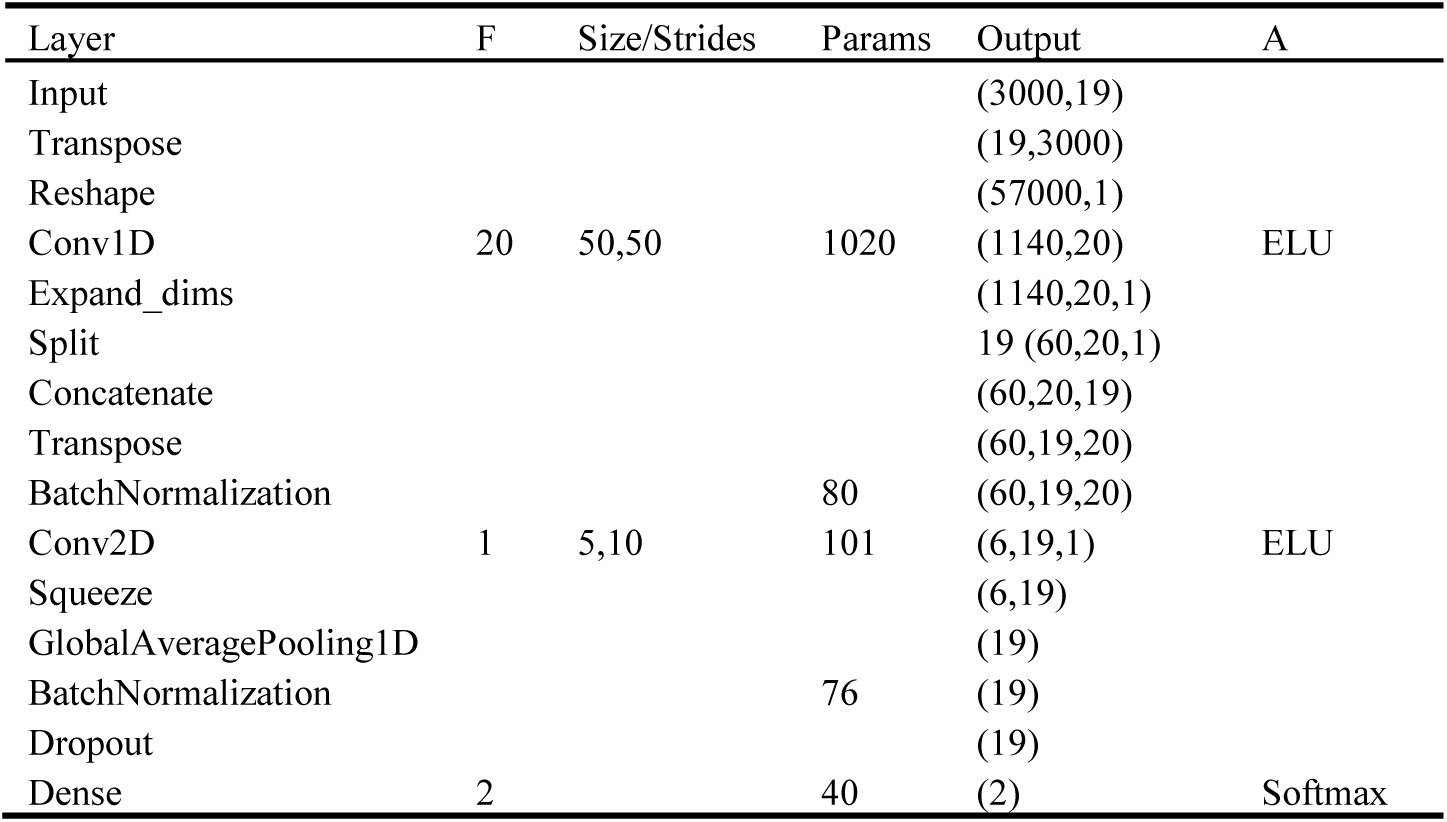
M3 Architecture, where F = number of filters or nodes, A = Activations. All conv layers use padding=’valid’, the he_normal initializer, and a max norm kernel constraint with a maximum value of 1. A learning rate of 0.001 was used. The total number of parameters is 1,317, with 1,239 trainable parameters. Layer names are from the Keras package [61].

#### Model Training Procedure

Our training procedure had 2 stages. (1) Hyperparameters were optimized across 10 training folds with the Hyperband algorithm in Keras-Tuner [62]. (2) After stage 1, training was repeated for 50 folds with the optimal hyperparameters. In both stages, 70%, 20%, and 10% of the data were used for training, validation, and test sets, respectively. To help ensure the cross-subject generalizability of our results, folds were generated with a group shuffle split approach that prevented samples from a participant being simultaneously mixed across training, validation, and test sets. When optimizing with Hyperband in stage 1, 75 initial epochs and a maximum of 200 epochs were used. A maximum of 200 epochs was also used when training in stage 2. To reduce training time, early stopping was employed if 20 epochs passed without a corresponding increase in validation accuracy (ACC) in both stages, and for stage 2, a checkpoint approach was used to retain model parameters from the epoch with maximal validation ACC. Across both stages, a class-weighted categorical cross-entropy loss function was employed, and a batch size of 128 with shuffling was used. We quantified model performance with ACC, balanced ACC (BACC), sensitivity, (SENS), and specificity (SPEC). To determine whether there were statistically significant differences in stage 2 model performance, we also performed a series of family-wise, paired, two-tailed t-tests between the ACC, BACC, SENS, and SPEC of each model followed by false discovery rate (FDR) correction applied separately to the p-values for each metric. The hyperparameters used when tuning with HyperBand can be found in our code repository.

### 2.3 Description of Explainability Analyses Applied to All Models

While the M2 and M3 architectures enable the application of a wider range of explainability methods than M1, we first compared the spatial and spectral representations learned by each of the models (i.e., their sensitivity to the loss of individual channels or frequency bands in the input data) with a pair of preexisting explainability analyses.

#### Spatial Explainability Analysis

To identify the sensitivity of each model to each EEG channel (a metric giving insight into the relative importance of each channel to the model), we used an ablation-based explainability approach [63]. Specifically, we calculated the ACC of each model for each of the 50 stage 2 folds. We then iteratively replaced each channel in the test data with values of zero and calculated the percent change in model performance after ablation (Equation 1). While it would have been ideal to use a line noise-related ablation approach [43] to reduce the likelihood of the creation of out-of-distribution samples [39], the MDD dataset that we used is a publicly available dataset that was uploaded with prior low pass filtering below the line noise frequency. Regardless, “zero-out” ablation approaches are common in literature [2, 46].

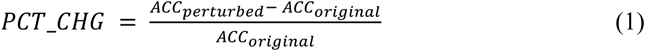

#### Spectral Explainability Analysis

To identify the sensitivity of each model to each EEG frequency band (i.e., the relative importance of each frequency band to the model performance, we used a perturbation approach [58, 63]. Specifically, we calculated the model test ACC for each fold, used an FFT to convert the test samples to the frequency domain, replaced the Fourier coefficients within a target frequency band with values of zero, converted the test samples back to the time domain with an inverse FFT, calculated the model test ACC on the perturbed data, and calculated the percent change in model ACC (Equation 1). We repeated this process for each canonical frequency band: δ (0-4 Hz), θ (4-8 Hz), α (8-12 Hz), β (12-25 Hz), γ (25-45 Hz). We used a zero-out approach here due to our relatively large number of folds and the computational intensiveness of many existing approaches that involve many repeated permutations across samples [13] or replacements of Fourier coefficients from a Gaussian distribution [11].

### 2.4 Description of Approach for Characterization of M2 and M3 Filters

Our previous explainability analyses were model-agnostic and enabled the comparison of models M1 through M3. However, in subsequent sections, we want to perform a series of model-specific explainability analyses that are uniquely enabled by our M2 and M3 architectures. The first stage in our series of novel explainability analyses involved characterizing the features learned by M2 and M3 in their first convolutional layer. Additionally, whereas previous model visualization-based studies have mainly focused on analyses of a single fold [4–6] or select number of samples [3], we adapted our analysis to find the overlapping patterns uncovered across all 50 training folds. To that end, we extracted the 20 first-layer filters (each 50 time points long) for each of the 50 folds, obtaining 1,000 filters (i.e., 50 folds x 20 filters). We then calculated the spectral power in each filter, removed the zero-frequency component, applied k-means clustering with 100 initializations, and assigned the filters to 3 clusters. After clustering, we visualized the cluster centroids for insight into the patterns extracted in each cluster. We clustered filters for M2 and M3 separately but afterwards manually aligned centroid labels for easier comparison.

### 2.5 Description of Novel Activation Explainability Analyses for M2 and M3

While our characterization of the filters gave insight into the spectral features extracted by the model, it did not provide insights into the multi-channel nature of M2 and M3. As such, we next performed a set of analyses that identified disorder-related differences in (1) per-channel cluster activations and (2) the correlations of channel activations.

#### Channel and Cluster-Level Activation Analysis

For insight into MDD-related aspects of the features extracted in each cluster of M2 and M3 filters across channels, we input the test samples into the models and extracted the first convolutional layer activations of each filter for each EEG channel. To enable cross-model comparison of activations, after extracting the first convolutional layer activations for all test samples for a given model, we separately normalized the activations of each sample by summing the total per-sample magnitude of the activations across all filters and dividing each sample activation by its corresponding total per-sample activation. From there, we calculated the activation mean along the time dimension for each filter and channel combination and calculated a second mean to obtain the mean activation for each cluster of filters in each channel. We next performed a series of two-tailed t-tests (α = 0.05) comparing the mean activations of MDD and HC samples for each channel and filter cluster before applying FDR correction. We separately analyzed the test sets of each fold and calculated the number of folds with positive versus negative t-statistics and post-FDR corrected significance for each channel and filter cluster.

#### Channel and Cluster-Level Activation Correlation Analysis

Inspired by inter-channel correlation analyses in EEG [21, 22], we next sought to determine whether M2 and M3 uncovered similar disorder-related patterns of EEG activity by examining the correlations of activations across channels. Specifically, we output activations for each test sample following the second convolutional layer. We then normalized activations separately for each channel and filter, summing the total activation for each channel and dividing the activations for each channel by their corresponding total activation. For each test sample, we next calculated the Pearson’s correlation over time between each a pair of channel activations (e.g., for 19 channels that resulted in a 19×19 correlation matrix). At that point, we performed a series of two-tailed t-tests between each correlation coefficient for MDDs and HCs, obtaining 171 p-values per fold to which we applied FDR correction. After identifying the correlations with significant MDD-related differences in each fold (corrected p < 0.05), we calculated the number of folds with significant differences.

## 3 Results and Discussion

In this section, we describe and discuss our analysis of model performance, differences in post hoc spatial and spectral explanations for M1 through M3, characterization of M2 and M3 first layer filters, disorder-related differences in spatial activations for each cluster of M2 and M3 filters, and disorder-related differences in M2 and M3 channel correlations. We lastly discuss some of the more important aspects of our approach, the implications of our architectures and post hoc explainability analyses for the broader field, limitations of our study, and opportunities for future work based upon our study.

### 3.1 M1-M3: Model Performance Analysis

Table 1 shows the mean and standard deviation of the performance of each of our models. All architectures performed well above chance-level. M1 had a much higher SENS than SPEC, while M2 and M3 had a greater balance between SENS and SPEC. Additionally, t-tests (p < 0.05) revealed that M1 had significantly higher ACC, BACC, and SENS than M2 and M3 following FDR correction. However, there was not a significant difference between M2 and M3 performance, which suggested that M3 was able to learn linearly separable features without dropping performance below that of M2. Many studies trained models that outperformed all three of our architectures for MDD classification [25, 31, 57, 64–66]. However, most of those studies used more traditional cross-validation approaches that allowed samples from the same recording to be distributed across training, validation, and test sets. As described in previous studies [20], this is a frequently occurring problem within EEG classification, as the heavy dependencies within EEG samples from the same sample result in data leakage between sets that prevents test performance from providing a realistic measure of model generalizability. Relative to most MDD classification studies on the same dataset that had valid cross-validation procedures, M1 performed comparably [17, 47, 58, 67, 68], though M2 and M3 lagged slightly behind. Nevertheless, we could potentially boost M2 and M3 performance with data augmentation [67] or transfer learning methods [58, 68] that have improved MDD classification performance in previous studies.

**Table 2.**
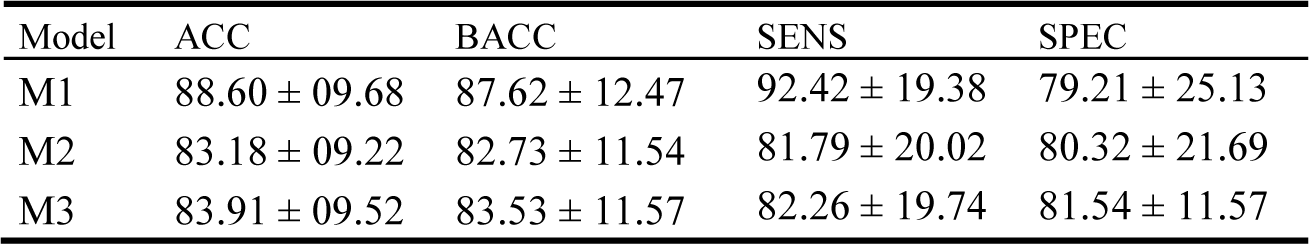
Model Performance Results.

| Model | ACC | BACC | SENS | SPEC |
| --- | --- | --- | --- | --- |
| M1 | $88.60 \pm 09.68$ | $87.62 \pm 12.47$ | $92.42 \pm 19.38$ | $79.21 \pm 25.13$ |
| M2 | $83.18 \pm 09.22$ | $82.73 \pm 11.54$ | $81.79 \pm 20.02$ | $80.32 \pm 21.69$ |
| M3 | $83.91 \pm 09.52$ | $83.53 \pm 11.57$ | $82.26 \pm 19.74$ | $81.54 \pm 11.57$ |

### 3.2 M1-M3: Post Hoc Explainability Analysis

Fig. 1 shows our post hoc spectral explainability analysis results. All models identified δ and β as having highest importance. However, M1 focused more on β, while M2 and M3 focused slightly more on δ. M1 did place some importance on α, and M2 and M3 did place some importance upon θ. These findings make sense within the context of existing literature. For example, the increased δ-band activity in MDD has often been described in previous studies [69] and has been suggested as a compensatory mechanism [70] that inhibits sensory networks that negatively impact attention and concentration in individuals with MDD [71]. Additionally, previous studies have found that MDD is characterized by abnormally high β power relative to lower frequency bands (δ, θ, α) [72] and have suggested that this increase in β is a compensatory mechanism that reduces impairment of attention, working memory, and executive processing [73].

**Fig. 1.**
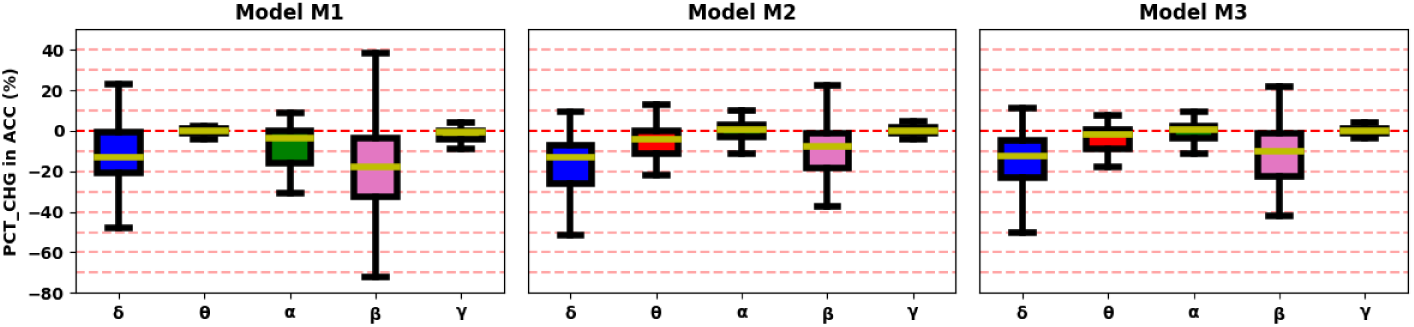
Post Hoc Spectral Explainability Results for Models M1 through M3. The leftmost, middle, and rightmost panels show the explanations for models M1, M2, and M3, respectively. The x-axis shows each canonical frequency band, and the y-axis is shared by all 3 panels and shows the percent change in ACC following the perturbation of a frequency band across all 50 folds.

Fig. 2 shows our post hoc spatial explainability analysis results. M2 and M3 had virtually no sensitivity to the loss of any one channel. That made spatial explainability more challenging but also provided a robustness that could be helpful in clinical settings [54] where data from key channels might be periodically lost, resulting in detrimental effects upon model performance (e.g., loss of Cz in M1 could lead to a 25% decrease in ACC). This reduced sensitivity to channel loss is atypical amongst deep learning models trained on raw EEG data [12, 54, 68], and potentially a unique and unexpected benefit of our architecture. It possibly results from how M2 and M3 consolidate all of the information from all 19 channels into 19 extracted features. For the models to perform well, they would need to efficiently extract the most important information in each channel and develop robust cross-channel representations. M1 does provide some helpful insights. Key M1 electrodes included Cz, P4, Pz, C4, O1, O2, F4, and F7. This finding is relatively consistent with previous studies [74] that have identified increased regional homogeneity (i.e., a measure of synchrony) in the right middle frontal gyrus (F4, C4) and right superior occipital gyrus (O2) and decreased regional homogeneity in the left superior occipital gyrus (O1). It is also consistent with previous studies [75] that have identified decreased fractional anisotropy (i.e., a measure of white matter density) in the left frontal lobe (F7), right frontal lobe (F4), right inferior fronto-occipital fasciculus (F4, C4, P4, O2), and corpus callosum (Cz, Pz). Additionally, previous studies have identified strong interhemispheric asymmetry that could explain why the majority of the top electrodes were in the right hemisphere or were central electrodes [76].

**Fig. 2.**
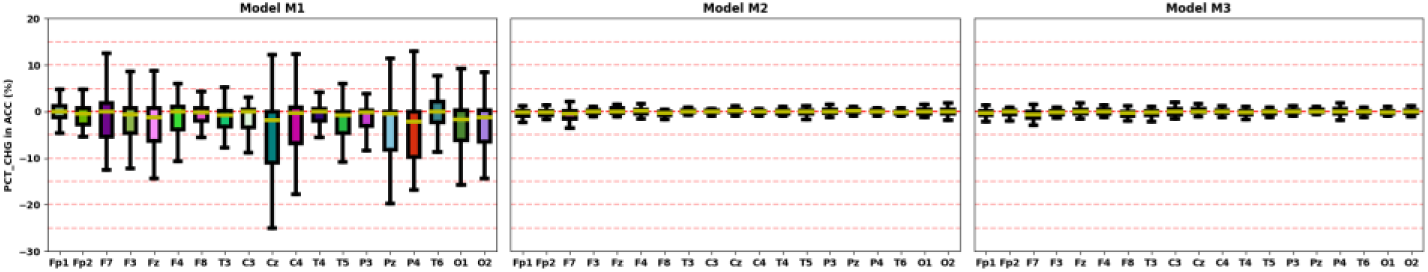
Post Hoc Spatial Explainability Results for Models M1 through M3. The leftmost, middle, and rightmost panels show the explanations for models M1, M2, and M3, respectively. The x-axis shows each channel, and the y-axis is shared by all 3 panels and shows the percent change in ACC following the perturbation of a channel across all 50 folds.

### 3.3 M2-M3: Characterization of Extracted Features

Fig. 3 shows the 3 M2 and M3 filter cluster centroids. Cluster 0 power was generally a bit unstable in lower frequency bands. It was characterized by lower δ at 2 Hz and higher δ at 4Hz, high to low α (from 10 to 12 Hz), and very low β power. Cluster 1 was a bit more stable, though it still had higher δ at 2 Hz and lower δ at 4Hz. Cluster 1 had high θ, moderate α and moderate β. Cluster 2 was similar to cluster 1 in having higher δ at 2 Hz and lower δ at 4Hz. It had particularly low values of θ and α but increased far above clusters 0 and 1 in β power. Clusters 0 and 2 tended to exhibit opposite behaviors, while cluster 1 tended to demonstrate more moderate behavior. Given these findings, one might expect the models to demonstrate higher cluster 2 activations in MDDs than HCs (i.e., due to increased β activity in MDDs [72]) and lower cluster 0 activations in MDDs than HCs (i.e., due to decreased β activity). The presence of consistently high to moderate cluster 1 δ might also result in higher activations in MDDs [69].

**Fig. 3.**
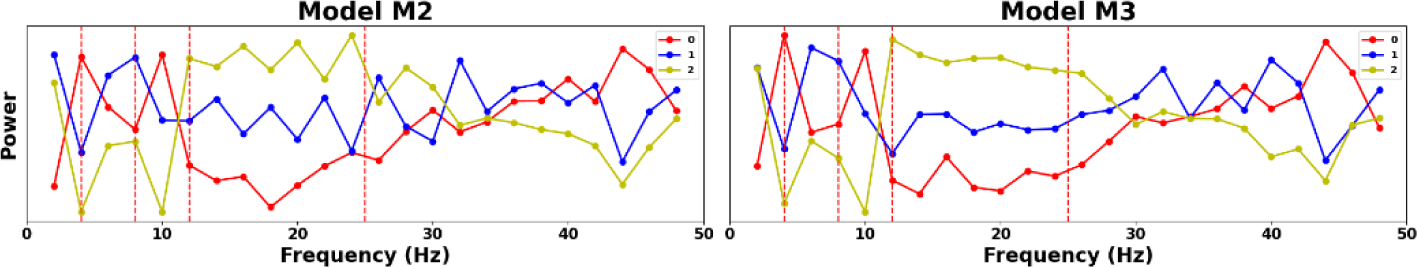
Filter Cluster Centroids for M2 and M3. The leftmost and rightmost panels show cluster centroids for M2 and M3, respectively. Cluster labels were aligned across models for easier comparison of centroids and subsequent analyses. Centroids 0, 1, and 2 are shown in red, blue, and yellow, respectively. Actual spectral power values are represented as points. Lines are included to make it easier to relate consecutive power values across frequencies. However, lines should not be considered representative of the actual trend of power values between consecutive points.

### 3.4 M2-M3: Post Hoc Spatial Activation Analysis

Fig. 4 shows our post hoc analysis of the mean first convolutional layer activations of each cluster of filters in each channel. Cluster 0 had higher activations in HCs than MDDs in both M2 and M3 across a relatively large number of folds (30-40%), and cluster 2 had higher activations in MDDs than HCs in both M2 and M3 in a number of folds (20-40%). Cluster 1 did have higher activations in MDDs than HCs in a small percentage of folds (10-20%). M2 cluster 2 had the greatest number of folds with significantly higher MDD activations in F4 and T3 (i.e., 26-28%), and M3 cluster 2 was highest in F3, Fz, F8, T3, and T4 (i.e., 34-36%). The higher M1 and M2 cluster 2 frontal activations could indicate that MDDs often had higher frontal beta activity, which fits with our spectral explainability findings and with existing literature on the compensatory nature of frontal β in MDD [72, 73]. Moreover, our cluster 2 findings fit with our cluster 0 findings, as HCs tended to have higher activations for cluster 0 (which was characterized by low β power) than MDDs. Lastly, M2 cluster 1 had higher activations in MDDs than HCs in a handful of models. That could be attributable to the consistently moderate to high δ activity within that cluster, as higher frontal δ activity has previously been associated with MDD [69, 70]. Nevertheless, M2 cluster 1 activations were overall not well represented across folds, so while frontal δ has been associated with MDD, we cannot make a strong case for increased frontal δ power in MDDs based on our results.

**Fig. 4.**
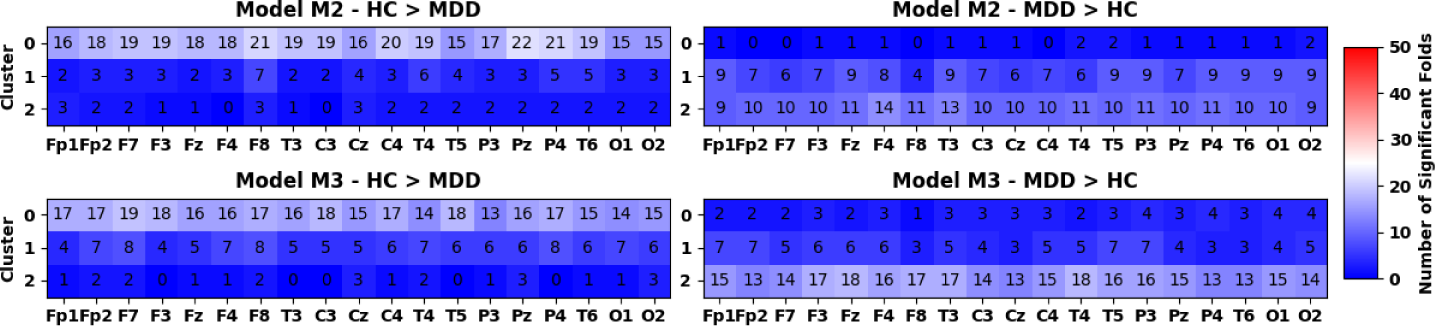
Disorder-Related Differences in Per-Channel Cluster Activation. The topmost and bottommost panels show results for M2 and M3, respectively. The leftmost and rightmost panels show the number of folds for which HCs had greater activation than MDDs (p < 0.05) in each cluster and the number of folds for which MDDs had greater activation than HCs (p< 0.05) in each cluster, respectively. All panels share the same color bar on the far right, with blue and red indicating significant differences in less than and greater than 25 folds, respectively.

### 3.5 M2-M3: Post Hoc of Activation Correlation Analysis

Fig. 5 shows the results of our post hoc analysis of the correlation between second convolutional layer activations for each channel. For ease of analysis, we focused on those correlations identified in more than 80% of the folds. However, there were other correlations present in a similar number of folds that did not exceed that threshold, and consistent with existing literature [77], MDDs generally had a higher degree of correlation between the channel activations across most channels. M2 and M3 identified 14 and 13 correlations present in more than 80% of models, respectively, with 8 of those correlations present across both models. Both models identified significant frontal-frontal correlations (M2/M3: Fz/F4; M2: Fp2/Fz, Fp2/F8; M3: F3/F4), frontal-central (M2/M3: Fz/Cz, Fz/C4; M3: F4/C4), frontal-parietal (M2: Fp2/Pz, F3/P4; M3: F4/P3), central-temporal (M2/M3: C4/T6), central-parietal (M2/M3: Cz/Pz, Cz/P4, C4/P4), and parietal-occipital (M2/M3: Pz/O2). Correlations overlapping across models included Fz/F4, Fz/Cz, Fz/C4, C4/T6, Cz/Pz, Cz/P4, C4/P4, and Pz/O2. The channels in which MDDs had the largest number of folds with greater activation correlation than HCs were mostly between frontal, central and parietal areas. Importantly, all of our largest overlapping correlations were within the right hemisphere or between right hemisphere and central electrodes. This could explain why M1 primarily identified right hemisphere and central electrodes as important to model performance. It also fits with previous studies that have identified the right hemisphere as particularly overactive in MDD [78], as having increased regional homogeneity [74], as having increased frontal (M2) and fronto-parietal (M3) β-band coherence [79], and as having increased frontoparietal connectivity (M3) related to reduced working memory [80].

**Fig. 5.**
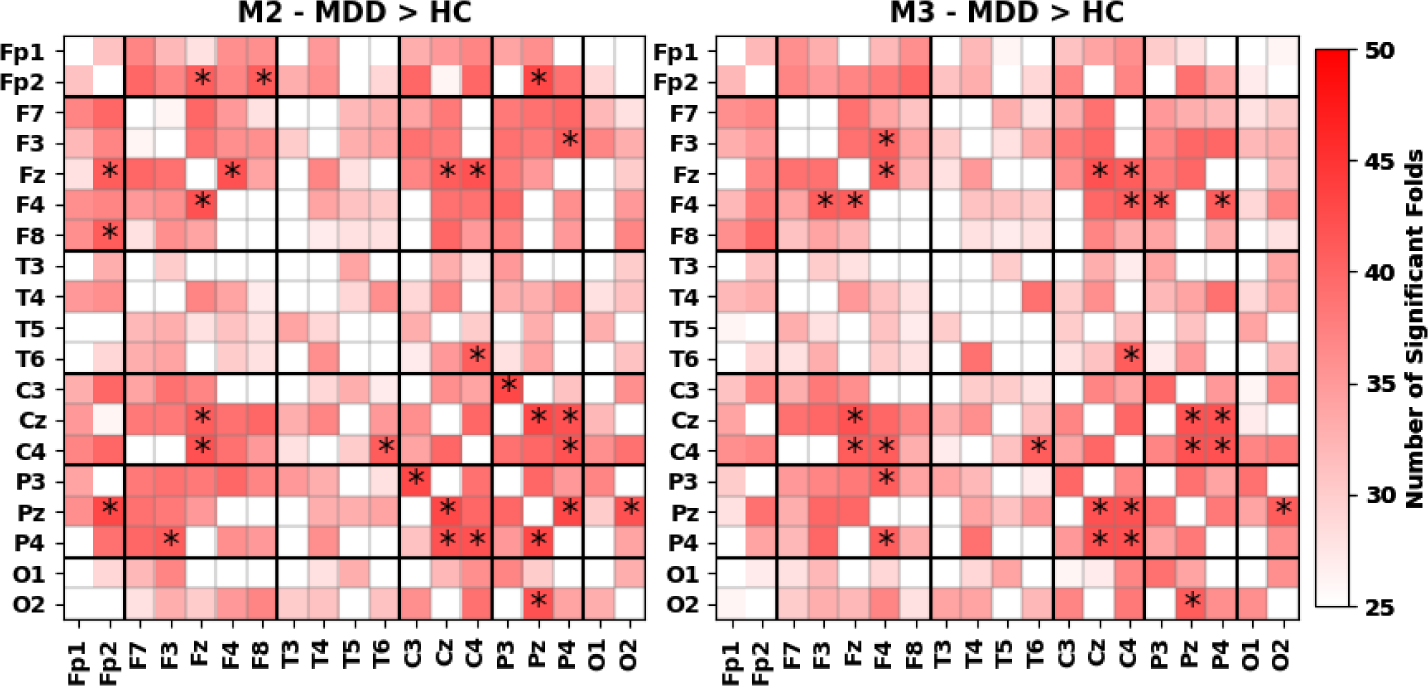
Disorder-Related Differences in Channel Activation Correlations. The leftmost and right-most panels show results for M2 and M3, respectively. Both panels show the number of folds in which MDDs had greater channel activation correlations than HCs. Results were displayed only for those channel correlations with significant disorder-related differences in more than 25 folds (i.e., 50% of folds), and asterisks were included for those channel correlations with significant differences in more than 40 folds (i.e., 80% of folds). Because HCs had only a couple channel activation correlations that were greater than MDDs in more than 25 folds, we did not include those results. Both panels share the color bar to the right. Channels are displayed on the x-and y-axes, and electrode groups (e.g., prefrontal, frontal, etc.) are separated by black lines.

### 3.6 Summary of MDD-Related Findings

In our perturbation-based spectral explainability analysis, all three models identified the β and δ frequency bands, which have often been related with MDD symptoms [71, 72], as most important. These findings are complemented our characterization of extracted features and spatial activation analysis for M2 and M3. Importantly, MDDs demonstrated higher activations across channels for filters with high β power (cluster 2), while HCs demonstrated higher activations across channels for filters with lower β power (cluster 0). Thus, the model often relied upon increased β power for the identification of MDDs. Increased β power in MDD is a well-characterized phenomenon and is likely a compensatory mechanism that alleviates attention, working memory, and executive processing impairment [72, 73]. Furthermore, MDDs had slightly higher activation for a cluster (cluster 1) with a higher average δ power, which indicates that our model likely relied on increased δ power in a few folds. Increased δ power in MDD is well-characterized [69] and is thought to be a compensatory mechanism that aids attention and concentration in MDD [70, 71]. Our spatial analysis of M1 primarily identified right hemisphere electrodes as most important, which fits with our channel activation correlation analysis results. We identified higher brain-wide inter-channel activation correlations in MDDs, with the strongest increases found mostly within the right hemisphere. This is strongly corroborated by studies that have identified higher right hemisphere activation [78] and synchrony [74] in MDD, particularly within the β-band [79].

### 3.7 Further Discussion on Approach Development

There are multiple key aspects of our approach that bear further discussion. (1) The first key aspect is our choice of long first convolutional layer filters. We chose a filter length of 50 so that the filter length would roughly equal half of a second of signal. While some previous studies have used longer filter lengths that do further enhance explainability [4, 5], having too long of a filter can increase the number of model parameters and decrease the flexibility of the representations that the model can learn. With a filter length of 50 time points at a sampling rate of 100 Hz, the filter length is long enough for an FFT that yields multiple frequency coefficients in the lower frequency bands (e.g., δ) that are associated with MDD. (2) The second key aspect of our approach is our choice to integrate activations within each channel. This enabled an additional level of spatial explainability wherein we could analyze MDD-related differences in the mean activations for each channel and the correlation between activations for each channel. It also built a bottleneck into the model that forced the model to extract the most salient representations within each channel. (3) The third key aspect of our approach was our choice to use comparable M2 and M3 architectures with the main difference being that M3 had only a linear classifier. This helped us evaluate whether our architecture could learn more interpretable features that might be distilled to the point of linear separability. (4) The next key aspect of our approach was our choice to cluster the first convolutional layer filters. This has been done in a few previous model visualization-based studies [4–6], and it is a critical component of our analysis. Because like EEG data, the filters were non-stationary in nature, converting them to the frequency domain enabled cluster centroids to be visualized in a relatively straightforward manner to characterize features extracted by the model. Additionally, while analyzing the 20 filters in each model on a per-model basis would have provided some insights, it would have been difficult to account for so many filters and their effects on each channel in subsequent analyses, and this problem would have only compounded when analyzing filters from all 50 folds. Without the clustering approach, obtaining model visualization-based multi-model insights into the features learned by the first convolutional layer would have been infeasible. While we could have used a larger number of clusters, it was necessary to strike a balance between having a larger number of clusters that might optimally separate the filters and having a relatively small number of clusters that would make interpretation of findings more intelligible. Given that interpretation was the goal of this study, we used a number of clusters identified as optimal in previous studies [4, 5]. (4) We applied our analyses to the test data, rather than the training data, with the goal of identifying patterns that were generalizable across subjects. (6) Lastly, we integrated our model explanations across a large number of folds with the goal of obtaining generalizable insights. This was a highly important decision given how dependent explanations can be upon random data distributions, random model initializations, and minor variations in architecture [53]. Because of this dependence, explanations could be extremely faithful to the model but still be ungeneralizable [55].

### 3.8 Broader Implications of Approach

Our approach was developed within the context of existing literature. Most previous model visualization-based [4, 5, 16] and activation-based [7] studies involved single channel data and analysis of a single fold, without suggesting a mechanism by which they might be extended to multichannel EEG data or provide global insight into a large number of models simultaneously. This is similarly the case in the relatively few model visualization-based studies with multichannel EEG data [3], which have generally only provided insight into a single fold or a few samples. Our approach offers a way to consolidate explanations across a large number of models with the goal of identifying more reliable biomarkers of neuropsychiatric disorders. Additionally, our channel activation correlation analyses are a feature uniquely enabled by our architectures. While two previous studies have developed interaction explainability approaches [17, 47], our analysis is one of the earliest in the field. More broadly, our architectures and post hoc analyses have implications for future multichannel EEG deep learning applications. They could provide insights into models trained on EEG data from a variety of other disorders (e.g., schizophrenia [12, 59, 60], Parkinson’s disease [81, 82], Alzheimer’s disease [83, 84]), and combined with our multi-model analysis approach, they could eventually identify novel disorder-related EEG biomarkers, similar to how explainable machine learning approaches have led to the discovery of biomarkers in neuroinformatics [85] and other domains [86]. Moreover, while we do not explicitly visualize or perform a waveform analysis in this study [4, 5, 16], the filters closest to each cluster centroid can be visualized in the time domain with the goal of identifying unique waveforms associated with a particular disorder.

### 3.9 Limitations and Future Work

In this study, we integrated activations from the first layer convolutional filters within each channel, which enhances spatial explainability but could reduce model performance. Future versions of our model could potentially integrate activations across channels. Additionally, we focused primarily on activation-based analyses. Other model visualization studies have used model perturbation-based approaches for insight into the key frequency bands and waveforms extracted within each first layer convolutional filter [4, 5]. We could do something similar with our approach by selecting the filters closest to each cluster centroid and perturbing them in the time domain or by converting them to the frequency domain, perturbing a particular frequency band, and converting them back into the time domain for prediction. Previous activation analysis studies have also used linear regression to identify linear relationships between early model activations and final predictions [6]. Rather than performing t-tests comparing activations for each class, we could do something similar for enhanced insight into the patterns that the model used for classification. Additionally, as previously stated, model explanations can be highly dependent upon the model training, validation, and test set distribution, upon model initializations, and upon minor variations of the model architecture. In this study, we accounted for that by analyzing explanations for a relatively large number of folds. However, in future studies, we might also use multiple initializations per fold or vary model hyperparameters. In this study, we chose to have a first convolutional layer filter length of 50 points (i.e., 0.5 seconds). Previous model visualization studies have used filter lengths of as short as 0.25 seconds [3] and as long as 2 seconds [4–6]. While the optimal filter length is based on a balance of model performance and enhanced explainability, future studies might systematically study the filter length parameter. It could also be helpful to evaluate the performance of our architectures on other multichannel EEG datasets to determine whether their utility is broadly generalizable across beyond our dataset, and even if our architectures themselves do not generalize well beyond our dataset, the principles they used related model visualization-based explainability could be useful in other applications. Lastly, our architectures obtain high levels of performance with a number of parameters an order of magnitude below the baseline architecture. It could be beneficial to examine in future studies how the performance of our models varies relative to a baseline architecture when the amount of training data is varied. Data augmentation [67] and transfer learning approaches might also provide an avenue towards enhanced performance for our architectures while retaining high levels of explainability [58, 68]. It should also be noted that the generalizability of any insights from our approach are still limited by the similarity of our MDD and HC participants to the overall population. As such, training our models on a larger MDD dataset could also help lead to the identification of more generalizable EEG biomarkers.

## 4 Conclusion

Deep learning methods are increasingly being applied in raw EEG-based classification studies but present unique problems for explainability that must be addressed if these models are to be used in clinical or medical research contexts. Moreover, if these models are to be used in medical research contexts to identify novel EEG biomarkers, the inherent randomness of patterns uncovered by models must be accounted for by combining explanations across a large number of folds, initializations, or architectures. Model visualization-based explainability approaches, which involve structuring deep learning model architectures such that extracted features can be characterized, have emerged as having high potential for faithfully representing what models have learned. However, they are generally underexplored within the domain of multichannel EEG, and approaches to combine model visualization-based explanations across many training folds have not yet been developed. In this study, we present two novel architectures that enable model visualization-based insights into (1) spectral features learned by models, (2) how those spectral features are distributed across channels, and (3) how the model relies upon correlation between channel activations. We compare our architectures to a baseline architecture, obtaining slightly lower performance and finding that our architectures rely upon similar spectral features while being much more robust to channel loss. We then perform a series of novel explainability analyses and find that across 50 folds relative to healthy controls, individuals with MDD exhibit higher β power, potentially higher δ power, and higher whole-brain correlation that is most strongly represented within the right hemisphere. This study provides multiple key insights into MDD and represents a significant step forward for the domain of explainable deep learning applied to raw EEG. We hope that it will inspire future efforts that will eventually enable the development of explainable EEG deep learning models that can contribute both to clinical care and the discovery of novel EEG biomarkers.

## Acknowledgments

This study was funded by NIH grants R01MH123610 and R01MH118695 and NSF grant 2112455.

## Disclosure of Interests

The authors have no competing interest to declare.

## Notes

https://figshare.com/articles/dataset/EEG_Data_New/4244171

https://github.com/cae67/InterpretableConv

### Competing Interest Statement

The authors have declared no competing interest.

